# A lumenal interrupted helix in human sperm tail microtubules

**DOI:** 10.1101/088476

**Authors:** Davide Zabeo, John M. Heumann, Cindi L. Schwartz, Azusa Suzuki-Shinjo, Garry Morgan, Per O. Widlund, Johanna L. Höög

## Abstract

Eukaryotic flagella are complex cellular extensions involved in many human diseases gathered under the term ciliopathies. Currently, detailed insights on flagellar structure come from studies on protozoa. Here, cryo-electron tomography (cryo-ET) of intact human spermatozoon tails showed a variable number of microtubules in the singlet region. Inside their lumen, a novel left-handed interrupted helix which extends several micrometers at their plus ends was discovered. This structure was named Tail Axoneme Intra-Lumenal Spiral (TAILS) and binds directly to 11 protofilaments on the internal microtubule wall, coaxial with the surrounding microtubule lattice. It leaves a gap over the microtubule seam, which was directly visualized in both singlet and doublet microtubules. We suggest that TAILS may stabilize microtubules, enable rapid swimming or play a role in controlling the swimming direction of spermatozoa.

## Introduction

Cilia and flagella can be found on many animal, plant and protist cells. It is an important cellular structure that can either act as an antenna (Hilgendorf et al., 2016), receiving signals from the environment, or provide cellular motility, such as in sperm tails. Malfunctions of human flagella are known as ciliopathies, and diseases of the motile cilium specifically are called primary ciliary dyskinesia (PCD). Patients with PCD present variable symptoms, such as pulmonary disease, situs invertus and infertility in both males and females (Praveen et al., 2015; Afzelius et al., 1995).

The flagellum consists of a membrane-covered axoneme, a complex arrangement of nine doublet microtubules (dMTs) surrounding two singlet central pair microtubules (CPs), interlinked with a multitude of protein complexes (Fisch and Dupuis-Williams, 2011). The dMTs consist of one complete A-tubule, made of 13 protofilaments, and a B-tubule, with 10 protofilaments (Nicastro et al., 2011a). In a complete microtubule made of 13 protofilaments, each α and β tubulin subunit is laterally adjacent and longitudinally offset to another subunit of its same kind. The only exception occurs at the seam, where an α and a β tubulin subunit are laterally in contact (Mandelkow et al., 1986; Kikkawa et al., 1994; McIntosh et al., 2009). This feature is part of the common lattice of each single microtubule. It is however currently unclear where the seam is located in axonemal microtubules. The location of this seam in the A-tubule has been suggested to be both at the dMT inner junction (Song and Mandelkow, 1995) and outer junction (Maheshwari et al., 2015; Ichikawa et al., 2017). Close to the distal tip of the eukaryotic flagellum, the B-tubule of dMTs often terminates and the A-tubule continues on, forming the so called “singlet zone” (Ringo, 1967; Satir, 1968). However, the presence of this singlet zone and its extent vary greatly (Höög et al., 2014). In rodent sperm, Woolley and Nickels (1985) observed that both the A-tubule and the B-tubule transition into singlet microtubules at the flagellum tip, coining the term “duplex microtubules” (Woolley and Nickels, 1985).

Microtubules are regulated by hundreds of microtubule-associated proteins (MAPs) and motor proteins (Nogales and Zhang, 2016; Howard and Hyman, 2007; Akhmanova and Steinmetz, 2015). MAPs fall into several categories based on their functions but, with regards to localization, they are classified into three main groups: those that bind the more dynamic plus end (+TIPs), e.g. EB1, XMAP215 and Clip170 (Perez et al., 1999; Mimori-Kiyosue et al., 2000; Nakaseko et al., 2001), those that bind and often stabilize the minus end, e.g. gamma tubulin and patronin (Goodwin and Vale, 2010; Kollman et al., 2010; Oakley and Oakley, 1989), and those that more generally interact with the microtubule outer surface, e.g. PRC1 (Chan et al., 1999; Kellogg et al., 2016). Motor proteins can also be found at all three locations, depending on the polarity of their movement (Scheffler et al., 2015). Altogether, in any given cell, the microtubule outer surfaces, plus and minus ends are often occupied by a wide assortment of MAPs and motor proteins that affect their characteristics, likely exerting effects in synergistic ways (Teng et al., 2001; Zanic et al., 2013).

Only two known MAPs, tau and tubulin acetylase, have been suggested to localize in the microtubule interior (Kar et al., 2003; Soppina et al., 2012). Yet, electron microscopy and (cryo) electron tomography have shown that proteins also localize to the inside of microtubules (Nicastro, 2006; Schwartz et al., 2012; Dentler, 1980; Rodríguez Echandía et al., 1968; Vaughan et al., 2006; Höög et al., 2007; Garvalov et al., 2006; Sui and Downing, 2006; Brown et al., 2016). These microtubule inner proteins (MIPs) are often found in dMTs and are all of unknown protein identity and function. However, their specific localization and variable frequency suggests that they serve important regulatory functions for the microtubule cytoskeleton (Nicastro et al., 2011b).

In this study, we performed the first three-dimensional reconstruction of intact human flagella using cryo-electron tomography on spermatozoon tails. By visualizing this cell type, we discovered a novel protein complex, Tail Axoneme Intra-Lumenal Spiral (TAILS), spanning several micrometers in the lumens of all healthy human spermatozoa axonemal microtubules at the distal end. TAILS binds to the inside of the microtubule and leaves a gap over the seam in single microtubules. The sub-tomogram average achieved sufficient resolution to resolve the tubulin subunits and the microtubule seam was directly visualized.

## Materials and methods

### Sample collection and plunge freezing

Spermatozoa were donated by three healthy men and frozen unperturbed in seminal fluid to which colloidal gold was added (to be used as fiducial markers). A Vitrobot climate-controlled plunge freezer (FEI Company Ltd., Eindhoven, The Netherlands) was used within 1-3 h post ejaculation. After freezing for electron microscopy, remaining cells were examined under the light microscope, where their motility ensured that viable spermatozoa had been frozen.

### Cryo-electron microscopy and tomography

Microscopy was performed as described previously (Höög et al., 2012; Höög and Lötvall, 2015). In brief, images (electron dose of ∼25 e^-^/Å^2^; -4 to -6 μm defocus) were acquired at 27500x on a Tecnai F30 electron microscope (FEI Company Ltd) operated at 300 kV. The detector was a GATAN UltraCam, lens-coupled, 4 K CCD camera (binned by 2) attached to a Tridiem Gatan Image Filter (GIF: operated at zero-loss mode with an energy window of 20 eV; Gatan Inc., Pleasanton, CA, USA). For tomography, tilt-series were acquired every 1.5 degree (±60 degrees) using serialEM software (Mastronarde, 2005). The total electron dose was kept between 80-120 e^-^/Å^2^. Cryo-ETs were calculated using eTomo and CTF correction was applied to all datasets.

### Subvolume Alignment and Averaging

Models with open contours approximating the path of microtubules in the electron tomograms were created in IMOD (Kremer et al., 1996). PEET (Nicastro, 2006) was used for adding model points with the desired spacing (8 nm, matching the periodicity of the TAILS complex) along each contour and for subsequent alignments and averages. For each model point a subvolume of 60 voxel^3^ (46.2 nm^3^) was selected. Number of tomograms, microtubules and particles for each average can be found in table 1. Subvolumes from individual microtubules were first aligned and averaged separately. Final alignment and averaging combining microtubules and tomograms was then performed starting from positions and orientations obtained by aligning the individual tube averages. Soft-edge cylindrical masks with empirically chosen radii were applied during alignment of singlet microtubules. Wedge-mask compensated principal component analysis followed by k-means clustering (Heumann et al., 2011) was used to check for heterogeneity and to assess the impact of missing wedge artifacts on candidate alignments.

**Table 1:**
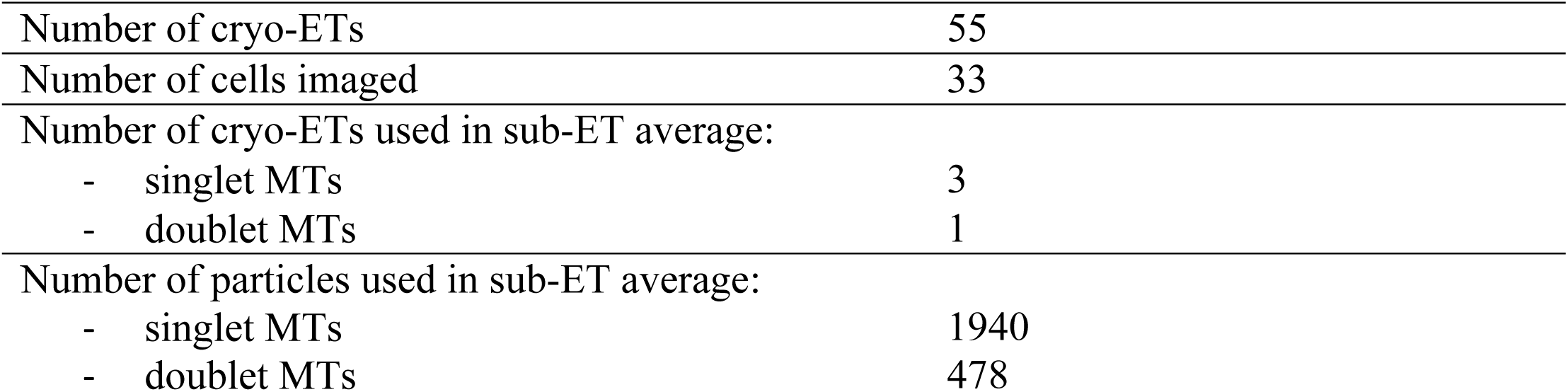
Number of electron tomograms acquired, cells imaged and sub-tomograms used for averaging.

### Statistical Analysis of the Singlet Region

Volumetric and isosurface representations corresponding to each tomogram were constructed by pasting the final average and its 3D model back into the original volumes. Statistical analysis of microtubule azimuthal positions was conducted in MATLAB (The Mathworks Inc., Natick, MA) using bootstrap analysis with 10,000 replicates and, when necessary, bounded depth-first search to compute the optimal value of the statistic of interest for each bootstrap replicate. Local measures such as mean and minimum neighbor angular distance or the number of neighbors closer than a specified angular distance showed little or no evidence of organization, with p-values ranging from 0.2 to 0.5. However, a more global measure shows strong evidence for higher-order organization. Specifically, we used bounded depth-first search to compute, for each spermatozoon, the length of the minimum angular distance path visiting each singlet microtubule exactly once. An overall probability of finding non-random orientation of microtubules based on data from all three spermatozoa was calculated. Combining the p-values for each spermatozoon using Fisher’s method for independent hypothesis tests (Fischer, 1925; Frederick and Fischer, 1948) showed that microtubules are non-randomly organized (overall p-value = 0.005).

Alternatively, we can also arrive at an overall p-value by bootstrapping the mean or, equivalently, the sum of the 3 minimum path lengths. Doing so resulted in only 24 successes out of 10,000 trails, corresponding to an even lower p-value estimate of 0.002. Both estimates are well below the conventional threshold of 0.05.

## Results

### Observation of an interrupted helix in the singlet microtubule lumen

To study the structure of human flagella, we performed cryo-electron microscopy of the distal tip of human spermatozoa that were plunge frozen in complete seminal fluid and we generated 55 cryo-electron tomograms on a total of 33 intact human sperm tails (Table 1). The exact location along the sperm tails and area included in 22 of those tomograms could be identified using lower magnification images (Figure 1A).

**Figure 1:**
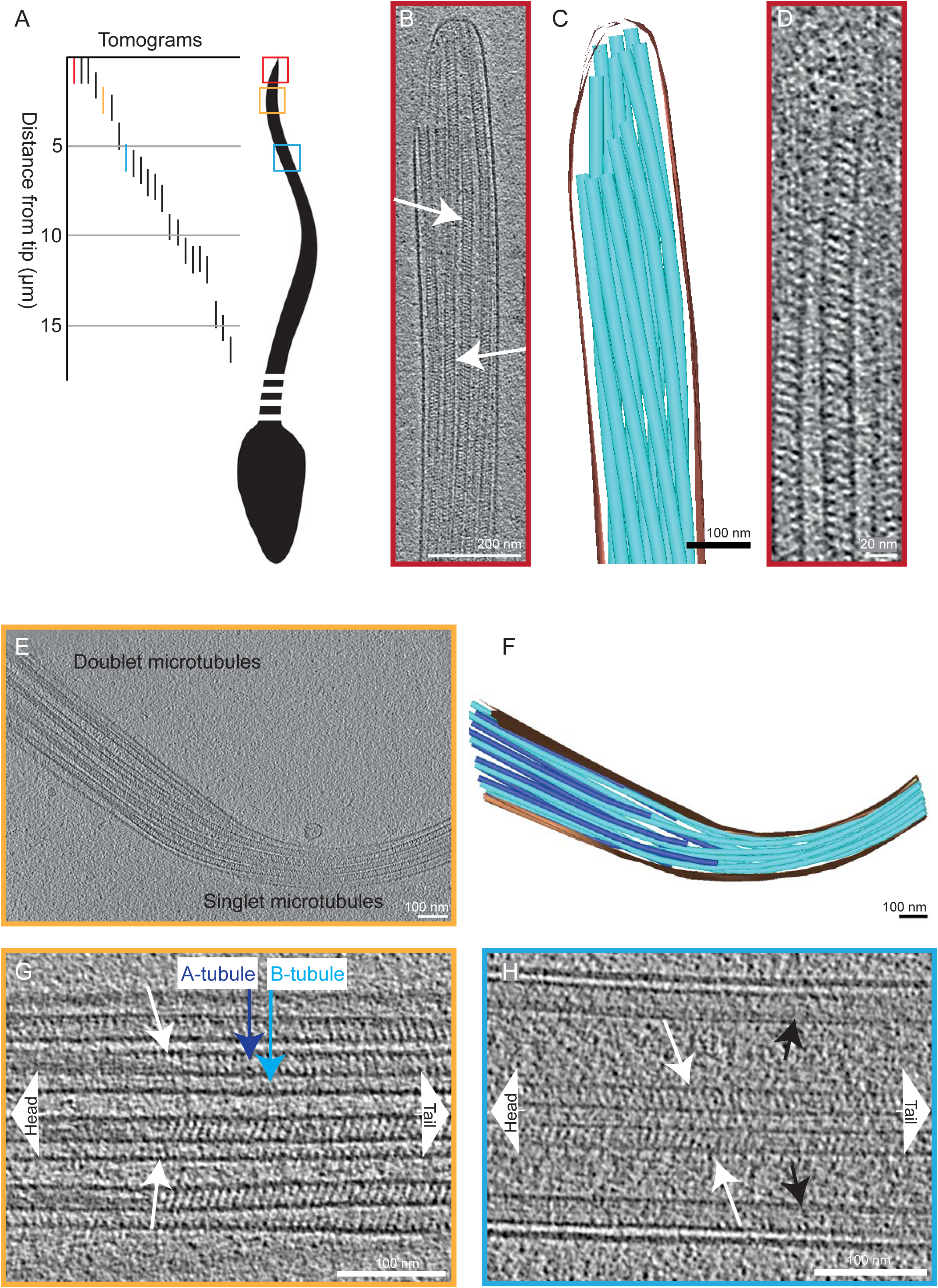
Microtubules in the end piece of the human spermatozoon show a repetitive pattern inside their lumen. A) A drawing illustrating positions along the sperm tails where cryoelectron tomograms were acquired. The 22 tomograms shown were acquired on a total of 13 different cells (1.7 tomograms/cell on average). The highlighted tomograms are shown in the panels with matching colors. B) A 15 nm thick slice from a cryo-electron tomogram of a sperm end piece showing singlet microtubules with repetitive diagonal striations with variable tilt directions (arrows). C) A 3D model of the sperm tip shown in panel B. The microtubules are shown in turquoise, the membrane in brown. D) A 7 nm thick tomogram slice of one singlet microtubule from the sperm end piece in B. E) A tomographic slice showing the transition area from doublet microtubules to singlet microtubules in the sperm tail. F) A 3D model of the sperm tail shown in panel E. The membrane is shown in brown, the A-tubules in turquoise and the B-tubules in blue. G) A tomographic slice showing the end of the intralumenal structure (white arrows) inside the doublet microtubule. H) Central pair microtubules also contain a repetitive structure inside their lumen (white arrows). Black arrows point at doublet microtubules.

The end piece of spermatozoa contained only singlet microtubules (Figure 1B-C). 23 cryo-electron micrographs of different human spermatozoa end pieces displayed an unexpected, extensive, and regular interior structure in all microtubules (Supplementary Figure 1). This structure was further analyzed and studied in the acquired tomograms (Figure 1B, D). Transverse lumenal slices of the electron tomograms offset from the center displayed diagonal electron densities with 8 nm periodicity. The tilt of the striations was reversed on opposing sides of the lumen (arrows in Figure 1B and Supplementary movie 1), suggesting a left-handed helix with a pitch of 8 nm. We named this structure Tail Axoneme Intra-Lumenal Spiral (TAILS).

To investigate the extent of the TAILS complex, cryo-electron tomograms further from the tip were examined. A transition from distal singlet microtubules into doublet microtubules was observed in one tomogram, approximately 2.5 μm away from the tip (Figure 1E-F). In this region, the TAILS structure was still continuous inside all the singlet microtubules and persisted into the doublet microtubules. TAILS eventually terminated approximately 300 nm into the dMT (Figure 1G). It thus extended for around 3 μm from the spermatozoon tip. A TAILS-like structure was also present in the lumen of the B tubules, starting from their plus end, and it extended, with its 8 nm periodicity, further towards the sperm head than TAILS in the A-tubule. Closer to the cell body, the A-tubule was filled with less defined electron density.

Intraluminal TAILS-like striations were also found inside central pair microtubules. The distance from the tip at which central pair microtubules terminated was variable between cells. The central pair microtubules stretched through one tomogram acquired at 5.5 μm from the sperm tip (Figure 1H). In another cryo-ET, acquired closer to the cell body, the central pair microtubules terminated at 7.5 μm from the tail tip. This indicates that central pair microtubules might extend less far than other axonemal microtubules. In both cryo-ETs the TAILS-like striations were found in the central pair microtubules (Supplementary Figure 2). In neither of these tomograms did the dMTs terminate nor contain TAILS, probably because this occurs closer to the tip, as described above. These results suggest a preference for TAILS to localize towards the microtubule plus ends.

**Figure 2:**
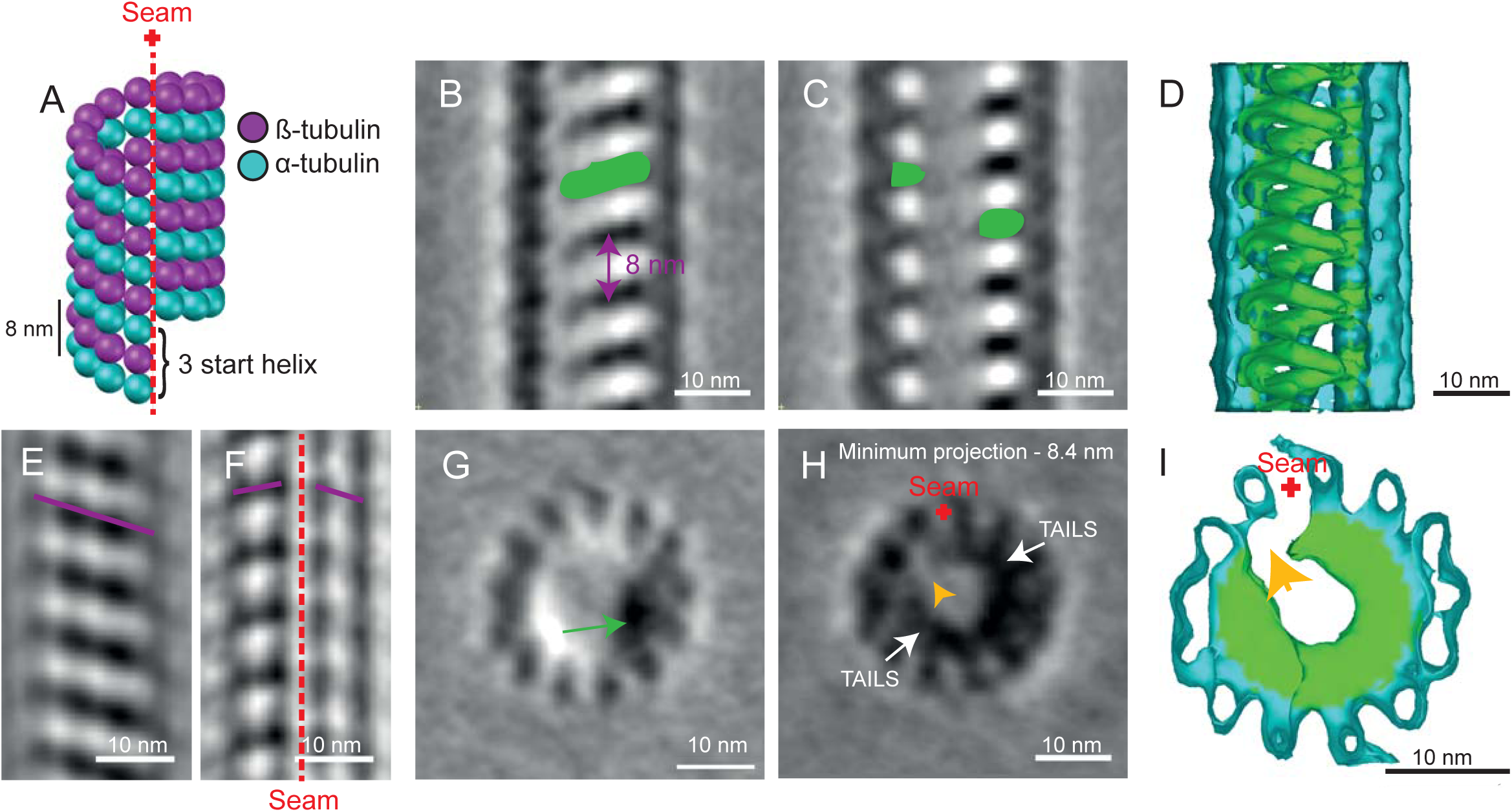
Sub-tomogram averaging reveals the TAILS complex, an interrupted left-handed helix that follows the pattern of the internal microtubule lattice. A) Cartoon of a typical 13 protofilament microtubule with the 3 start helix and a 12 nm pitch. The microtubule seam is marked with the red dotted line. B-C) 0.7 nm thick longitudinal slices through the sub-tomogram average showing the microtubule lattice and the internal electron density (green). D) The 3D model shows the microtubule lattice (turquoise) and the intralumenal structure (green). Some protofilaments have been cut away from the model to reveal the internal helical structure. E) A 4 nm thick slice of the sub-tomogram average shows a B-lattice arrangement of the tubulin heterodimers (purple line). F) A 4 nm thick slice of the sub-tomogram average shows a disruption in the microtubule B-lattice, revealing the location of the seam (red dotted line). G) A 0.7 nm thick cross-sectional view of the sub-tomogram average. The green arrow points at the electron density of TAILS. View from flagellum tip, looking towards sperm head. H) Projection of a 8.4 nm thick cross-sectional view of the sub-tomogram average showing the end of one TAILS complex segment, the beginning of the next and the gap between them (yellow arrow). I) 3D model of the microtubule (turquoise) and the TAILS complex (green) reveals the gap (yellow arrow) in the TAILS complex.

### The TAILS complex binds to 11 protofilaments with a gap over the seam

At first sight, it is difficult to see how one might reconcile an 8 nm pitch interior helix with the 12 nm pitch expected at the seam of the 13-protofilaments wall of singlet microtubules (Figure 2A). To resolve this discrepancy and clarify the nature of the structure, we performed sub-volume alignment and averaging on 1940 particles that were selected along 34 singlet microtubules.

Sub-volume averaging increased the signal-to-noise ratio revealing a left-handed interrupted helix. In longitudinal view, TAILS was clearly recognizable as the repetitive pattern seen in the raw data (Figure 2B-C; Supplemental Movie 2). A 3D model of the sub-tomogram average was created using density thresholding, which also showed the structure of TAILS (Figure 2D; Supplemental Movie 2). The average showed that TAILS is made up of multiple helical segments. One TAILS segment occurs every 8 nm along the microtubule axis. Each segment has a pitch of 12 nm, matching that of 13 the protofilament microtubule wall (Figure 2A). This led us to examine the site of TAILS binding in relation to the microtubule seam. The seam could be directly identified in the microtubule lattice (Figure 2E-F; Supplementary Figure 3).

**Figure 3:**
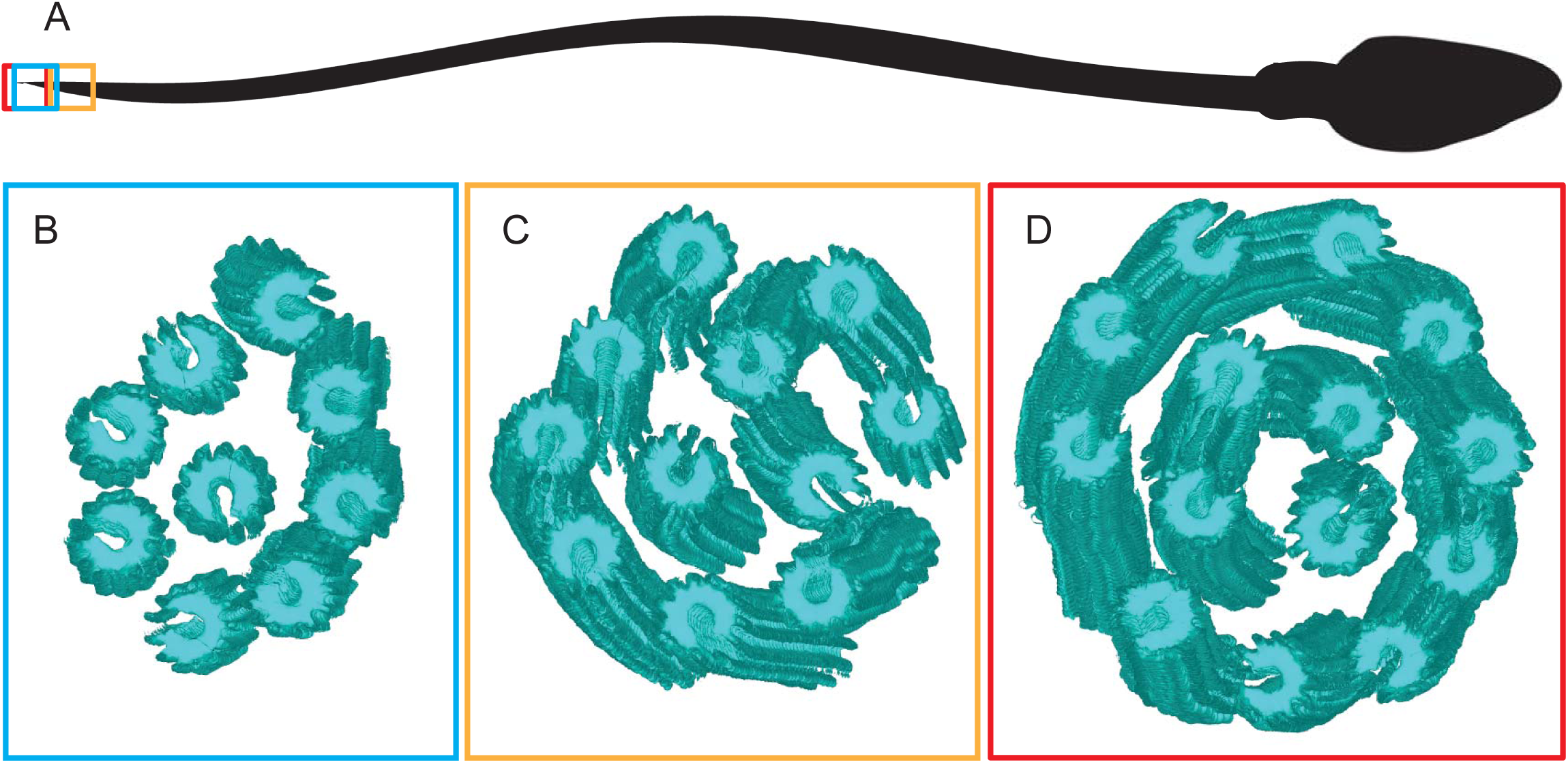
Number of microtubules in the singlet region vary but are non-randomly oriented. A) A cartoon of a spermatozoon showing the location of the cryoETs shown in B-D. Isosurfaces of sub-tomogram averaged singlet microtubules were aligned into a model volume according to their original orientations. The gap in the microtubules is used as a marker of the microtubule seam. Three singlet regions with A) 9 B) 11 and C) 14 microtubules were analyzed and showed a non-random orientation of the seam.

In cross-sectional view the TAILS complex is clearly visible as an electron density (Figure 2G) which rotates around the lumen when viewed at successive positions along the microtubule axis (Supplemental movie 3). Moving through the sub-tomogram average, gaps in the TAILS complexes are apparent on the inside of the microtubule (Figure 2H-I). Each segment spans between 240 and 305 degrees (based on centers or edges of electron densities, respectively) leaving a gap spanning portions of 2 protofilaments around the MT seam. The proximal end of a given segment occurs at approximately the same axial height as the distal end of the previous segment. The result is a stack of identically oriented, C-shaped segments that are coaxial with the surrounding tubulin helix. TAILS can also be thought of as an interrupted or incomplete 3 start helix, where the interruption is found across the microtubule seam.

### Microtubules in the human singlet zone are non-randomly organized

The singlet zone appears less highly organized than the more cell body proximal complete axoneme. In the human spermatozoa investigated here, the number and position of microtubules are variable both within the singlet zone of an individual single sperm tail, and between singlet zones of different sperm tails. The disruption of TAILS decoration over the MT seam can be used as a compass to investigate whether or not microtubule seams in the singlet region are randomly oriented with respect to each other.

Three cryo-ETs of singlet regions containing 9, 11 and 14 microtubules were investigated, two were acquired at the absolute tip of the sperm tail and one at 1.5 µm from the tip (Figure 3A). The presence of 14 singlet microtubules in one cryo-ET is noteworthy since it shows that both A-and B-tubules nucleate singlet microtubules. All of these microtubules consisted of 13 protofilaments, meaning that the incomplete B-tubule gained 3 protofilaments after separating from the A-tubule. Using the coordinates applied to rotate particles and generate the sub-tomogram average, the 3D isosurface models of singlet microtubules could be docked back into a 3D model representing the microtubule orientation inside each cell, allowing easy visualization of the seam orientations (Figure 3B-D).

To investigate possible non-randomness of singlet region seam orientations, bootstrap analysis was performed using a number of local and global measures of orientation. Local measures, such as mean or minimum angular distance to neighboring tubules, showed no indication of non-randomness. A more global measure, total change in angle for a path visiting each microtubule exactly once, does however suggest partial ordering. Bootstrapped p-value estimates of 0.011, 0.301, and 0.031 were obtained for singlet zones with 9, 11, and 14 microtubules, respectively. Combined data from all 3 singlet zones yields a highly significant p-value <= 0.005, as described in Materials and Methods, indicating that seam orientations are not completely random. We suggest that the observed deviations from randomness likely reflect remnants of the more rigid organization of the proximal axoneme.

### The seam is located between protofilaments A8 and A9 of the doublet microtubules

Since the position of the A-tubule seam in dMTs has not been resolved using cryo-ET, and other methods such as single particle analysis have given conflicting results (Song and Mandelkow, 1995; Ichikawa et al., 2017; Maheshwari et al., 2015), there is still some debate as to where it is localized. The orientation of the gap in TAILS can be used as an indicator of the microtubule seam position in dMTs.

In two cryo-ETs that we acquired, the transition from dMTs to singlets was included in the volume. The TAILS complex clearly extended into dMTs in one of these tomograms (Figure 1G). In the other cryo-ET, the TAILS complex in the A-tubule was not visible but B-tubules still had clear luminal decorations. This shows that the extent of the TAILS complex within dMTs is variable. Only the tomogram with complete TAILS decoration in both dMTs (478 particles) was used for sub-tomogram averaging. The structure has an 8 nm repeat, like in singlet microtubules, and a gap of the TAILS complex was found to be at the outer junction between protofilament A8 and A9 of the dMTs (Figure 4A-F; Movie S4). Thus, we inferred that this is the position of the A-tubule seam. Inside the B-tubule seven or eight protofilamens were decorated, possibly with a TAILS-like protein but protofilament B1, B10 and possibly B9 were undecorated.

**Figure 4:**
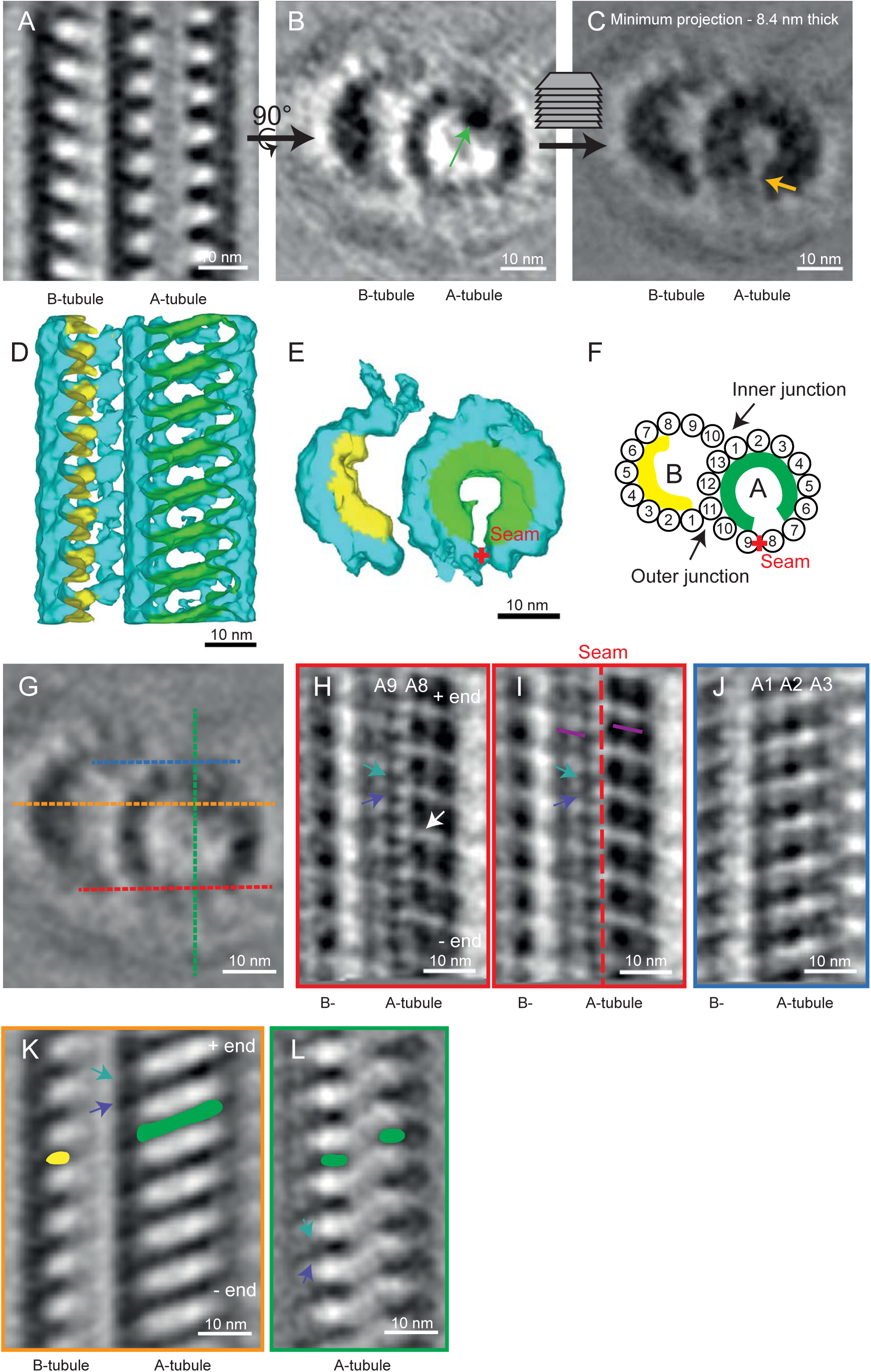
TAILS binds to the interphase between α and β tubulin in doublet microtubules. The A-tubule seam is found close to the outer junction, between protofilament A8 and A9. A) A longitudinal section through a sub-tomogram average of a dMT shows an 8 nm repeat structure on the inside of the microtubule walls. B) TAILS is seen as an electron dense dot in one cross-section of the average (green arrow). C) A minimum projection is done over an 8.4 nm thick cross-section which shows the expansion of TAILS over 12 protofilaments and the gap in the structure (yellow arrow). D) An isosurface model of the averaged dMT: tubulin in turquoise, the TAILS complex in green and the B-tubule decorations in yellow. The front microtubule wall has been cut away for better visualization. E) cross-sectional view of the isosurface from the plus end. The position of the seam is highlighted by the red cross. F) A cartoon of the dMT with protofilament numbers, the seam marked with red cross and TAILS extension in green. G) A cross-sectional view shows the location of the H, I, J and K longitudinal views. H) A 0.7 nm thick tangential slice of the average, showing the microtubule lattice from the outside. The resolution is below 4 nm since the subunits of the tubulin heterodimer were resolved. Due to the position of the MT plus end and the position of the gap (white arrow) in the lattice we infer the position of α (purple arrow) and β (turquoise arrow). I) A 4 nm slice of the average contains whole protofilaments and reveals the position of the A-tubule seam (red dotted line). The slope of the B-lattice is visualized on both sides of the seam with the purple lines. Because of a distance between the tubulin heterodimers we can infer the position of α and β tubulin. J) A 4 nm thick slice of the dMT sub-tomogram average, seen from the inner junction, shows a perfect B-lattice arrangement over protofilaments A1, A2, and A3. K) The inside of the microtubule average with the TAILS complex marked in green. TAILS binds between the α and β subunits. The decoration within the B tubule is marked in yellow. L) This view shows that TAILS binds to the interface between the α and β subunit.

Sub-tomogram averaging enabled to increase the signal to noise ratio to the point where the α and β tubulins are resolved in our dMTs (Figure 4G). The seam is clearly visible between protofilament A8 and A9 (Figure 4G-I), in the outer junction of the dMT. In comparison, the dMT inner junction shows a perfect B-lattice (Figure 4J).

Tubulin heterodimers are equally spaced 8 nm apart, but there appears to be a compaction of their electron densities that may or may not be TAILS related.

## Discussion

This is the first time an intact human flagellum has been studied using cryo-electron tomography. We have described a novel structure that we named TAILS, which is comprised of helical segments spaced every 8 nm. Many TAILS segments assemble into an array along several micrometers of the inner wall of all singlet microtubules in the end piece of human sperm tails. TAILS binds to the inside of the tubulin heterodimers and spans 11 protofilaments. The interruption of the helix spans over the position of the microtubule seam. TAILS has a pitch and handedness matching that of the tubulin helices comprising the microtubule wall. This pattern is consistent with what would be expected for a structure comprised of repeating identical segments or subunits capable of distinguishing between α and β tubulin. We suggest two alternative models for the molecular structure of TAILS, which could consist of multiple monomeric segments (Figure 5A) or multiple groups of 11 subunits (Figure 5B), leaving a gap over the seam in either case. Our sub-tomogram averaging yielded a resolution that was high enough to distinguish some of the tubulin subunits on the dMTs still inside of the frozen cell in its native state. The gaps between heterodimers have previously been shown to be larger than those between the subunits of each dimer (Alushin et al., 2014), therefore we consider it most likely that this gap in the microtubule lattice occurs in between heterodimers. We infer the identity of the tubulin subunits with the help following information: 1) microtubule plus end is capped with the β-tubulin subunit (Mitchison, 1993), and 2) flagella have their microtubule plus ends at the flagellum tip (Euteneuer and McIntosh, 1981; Song and Mandelkow, 1995). TAILS binds in the interface between the α and β tubulin subunits, and extends with a slight slope towards the microtubule plus end (Figure 4G and K-L).

**Figure 5:**
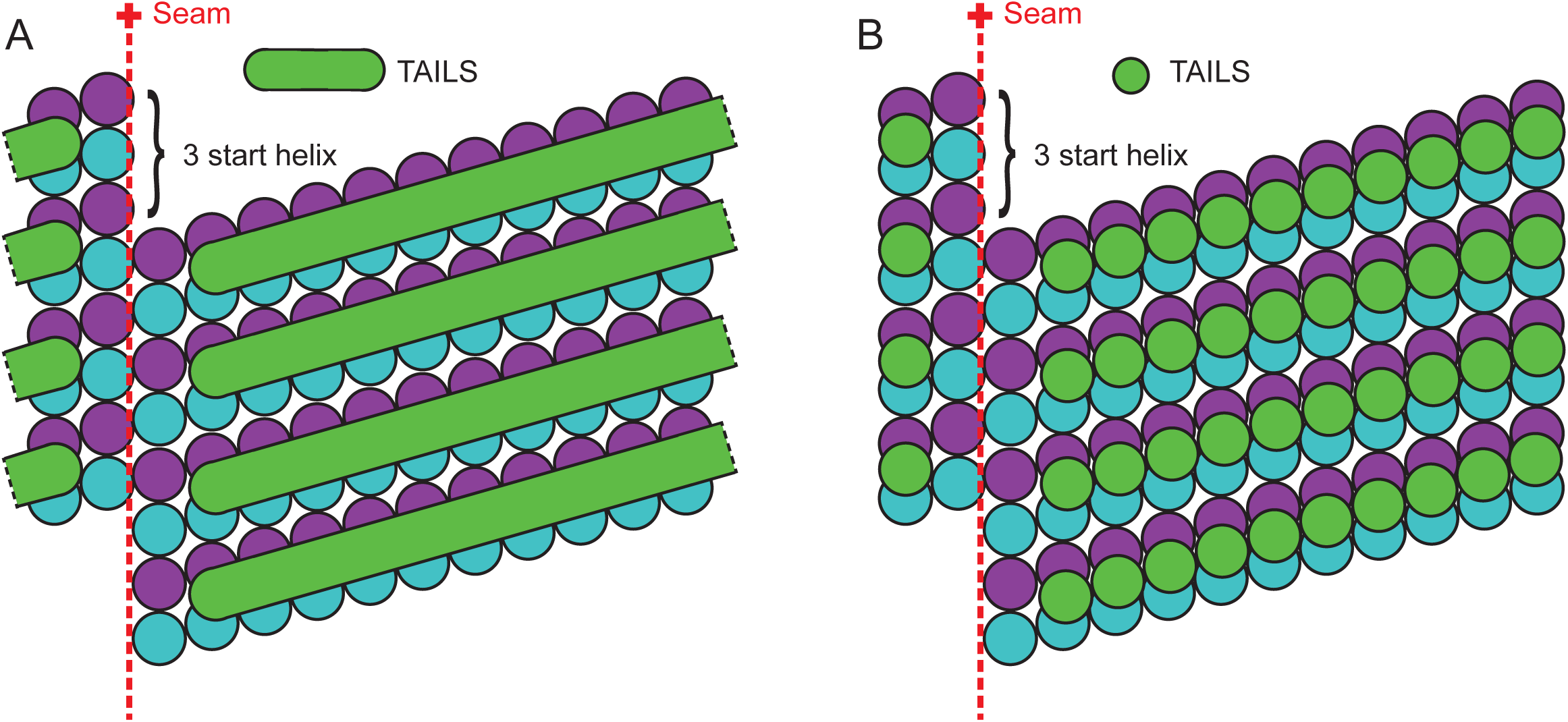
A schematic representation of two different models of the molecular structure of TAILS. A 13-protofilaments microtubule from the sperm tail tip is shown as opened up and unfolded into a sheet, viewed from the lumenal side. The TAILS complex (green) is placed at the position along the y-axis as found in dMTs, as we do not know the exact location in singlet microtubules. The α and β tubulin subunits are shown in blue and purple respectively. A) TAILS is drawn as monomeric C-shaped segments that bind to each protofilament at the intraheterodimer interface, leaving a gap over the seam. B) TAILS is drawn as a series of C-shaped multimeric segments, each formed by 11 subunits.

The achieved resolution allowed identification of the seam in both singlets and dMTs, making this the first time this structure has been visualized directly inside a cell. The reported position of the seam in dMTs differs with previously published papers. First, the seam was inferred to be at the inner junction of the dMTs of sea urchin sperm (*Psammechinus miliaris*) (Song and Mandelkow, 1995), then two more recent papers, using single particle techniques to achieve high resolution maps of dMTs from *Tetrahymena thermophila*, suggested a position of the seam at the outer junction, between protofilaments A9-10 (Ichikawa et al., 2017) or A10-11 (Maheshwari et al., 2015). We find the seam between the A8-9 protofilaments, both by the location of the gap in TAILS and by direct visualization. The differences observed might be a result of using different methods, or it might reflect a real difference in the location of the seam between species.

The presence of 14 microtubules in the singlet region shows that also in humans, the dMTs can split and form a duplex microtubule, like the ones seen in rodent spermatozoa (Woolley and Nickels, 1985). The 13 protofilament microtubule lattice found *in vivo* has been shown to be determined by the nucleation factor γ tubulin ring complex (γ-TuRC)(Moritz et al., 1995). That all the singlet microtubules consist of 13 protofilaments, even those derived from the 10 protofilaments B-tubule, argues that γ-TuRC is not the sole determinant of protofilament number *in vivo*. The orientation preference of the microtubules in the singlet zone suggests that the singlets maintain the seam position previously established in the complete axoneme.

TAILS decorates the inside of every tubulin heterodimer, except the two protofilaments at the seam, for several micrometers before the microtubule plus ends. The function of this complex structure is of course of highest interest. We suggest four different hypotheses, which are not mutually exclusive. First, TAILS seems to have compacted the tubulin heterodimer, a feature which would translocate to the outside of the microtubule. This rearrangement of the lattice might prevent motor proteins and other MAPs to bind in their regular way, defining the singlet region in a molecular manner. Second, TAILS might provide extra rigidity to the microtubules in this region, so that the end piece has an increased stiffness, which might aid motility, crucial to spermatozoa. It may provide a structural support such as the para-flagellar rod in *Trypanosoma brucei* (Höög et al., 2012; Portman and Gull, 2010; Koyfman et al., 2011), another single cell whose motility is crucial for its survival (Broadhead et al., 2006). Third, the TAILS complex could also stabilize microtubules, preventing the dynamic turnover constantly occurring in flagella tips (Marshall, 2001). Helical reinforcement is a low weight solution regularly used in engineering, *e.g.* in bicycle frame tubes or in armored hoses. Since microtubules depolymerize by outward curling of protofilaments (Simon and Salmon, 1990; Mandelkow et al., 1991), TAILS could prevent splaying similar to how spiral hose reinforcement prevents radial expansion. The saved energy could then instead be invested in rapid translocation. This hypothesis is consistent with the observation that the TAILS complexes extend to the central pair microtubules, far into the sperm tail. Lastly, TAILS might also play a role in determining the direction in which spermatozoa swim, since there is evidence in other organisms that the singlet zone may be associated with signaling or sensory functions as well as motility (Fisch and Dupuis-Williams, 2011).

Potential mechanisms for controlling the extent and orientation of the helical segments are worth considering. The TAILS complex has a gap in the structure spanning the inside of the microtubule seam. Binding specificity of the putative inner wall binding monomers coupled with conformational variation of the subunits closest to the seam alone could account for the observed structure. A monomer binding to the alpha and beta tubulin interface, like TAILS, could be unable to bind to the intraheterodimer interface across the seam, leaving a gap. If a bridge over this gap is present, such a structure would likely be comprised of protein(s) distinct from the inner wall binding monomers, and its length might naturally limit the span of the helical segments.

We performed cryo-electron microscopy and tomography on intact human spermatozoa. This study led to the finding of a novel, complex structure inside of the microtubule lumen, even though microtubules have been studied for over 50 years. This underlines the importance of studying human microtubules and flagella, as well as other organisms, by cryo-ET. Since sperm motility and morphology are determinants of male fertility (Esteves, 2016), understanding the functional role of the TAILS complex may have clinical implications relating to male infertility and contraception. We look forward to the future identification of the proteins involved in regulating the microtubule cytoskeleton from the lumenal side, including the proteins forming the TAILS complex.

## Acknowledgements

We thank R. Neutze for helpful discussions and for contributing funding from the Göran Gustafsson Foundation for Research in Natural Sciences and Medicine (awarded in Stockholm May 2016) and Wallenberg grant KAW2014.0275 and KAW2012.0106. We thank Dr. J. Van Blerkom at Colorado Reproductive Endocrinology for providing samples for our study, and J. R. McIntosh and A. Hoenger for helpful discussions. JHL was supported by a Sir Henry Wellcome Postdoctoral grant and a VR young investigator grant. Electron microscopy was done at Boulder EM Services Core Facility in MCDB, the University of Colorado.

## Author contribution

Experimental design: JLH, POW. Acquisition of cryo-electron micrographs: JLH, GM. Acquisition of cryo-electron tomograms: JLH, CLS. Calculation and modelling of tomograms: JLH, DZ. Sub-tomogram averaging and processing of data: JMH, DZ and ASS. Writing of manuscript: JMH, POW, DZ and JLH.

## Competing interests

Authors declare no competing interests.

## Supplementary material

**Figure S1:**
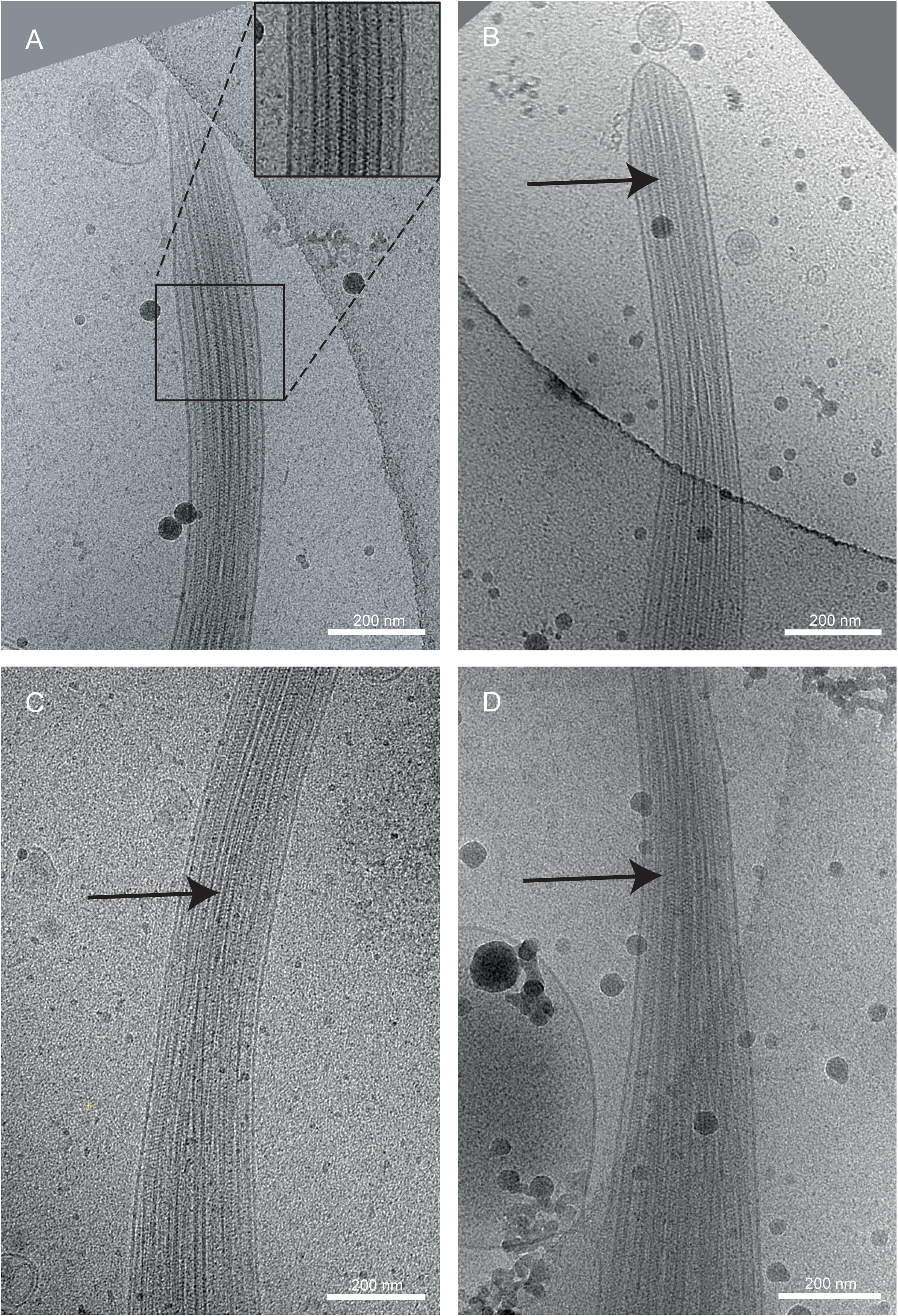
Cryo-EM of the end pieces of four intact human spermatozoa derived from three different donors. A) A cryo-electron micrograph of one sperm tail tip showing repetitive decoration of microtubules (zoomed image). B-D) In all sperm tips the repetitive pattern inside the microtubules in the singlet region is clearly visible. The arrows point to some of the more apparent places, but the helical pattern is present through most, if not all, visible areas.

**Figure S2:**
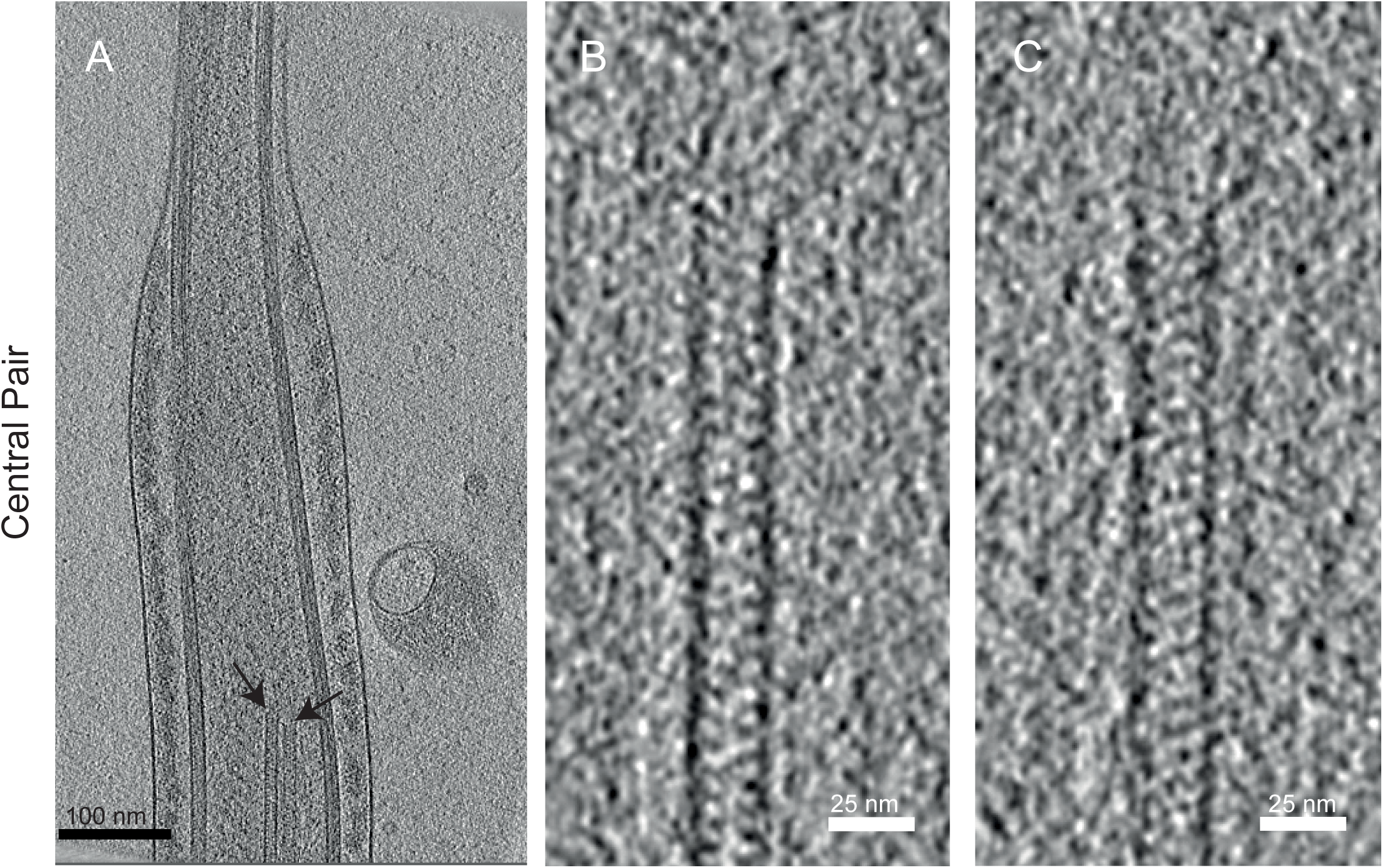
A TAILS-like complex is also present in the terminal parts of the central pair microtubules, around 7.5 micrometers from the sperm tip. A) Slice from a cryoelectron tomogram showing the distal ends of the central pair microtubules (black arrows). B-C) Zoomed-in images of the central pair microtubules show the striations resembling those typical of the TAILs complex.

**Figure S3:**
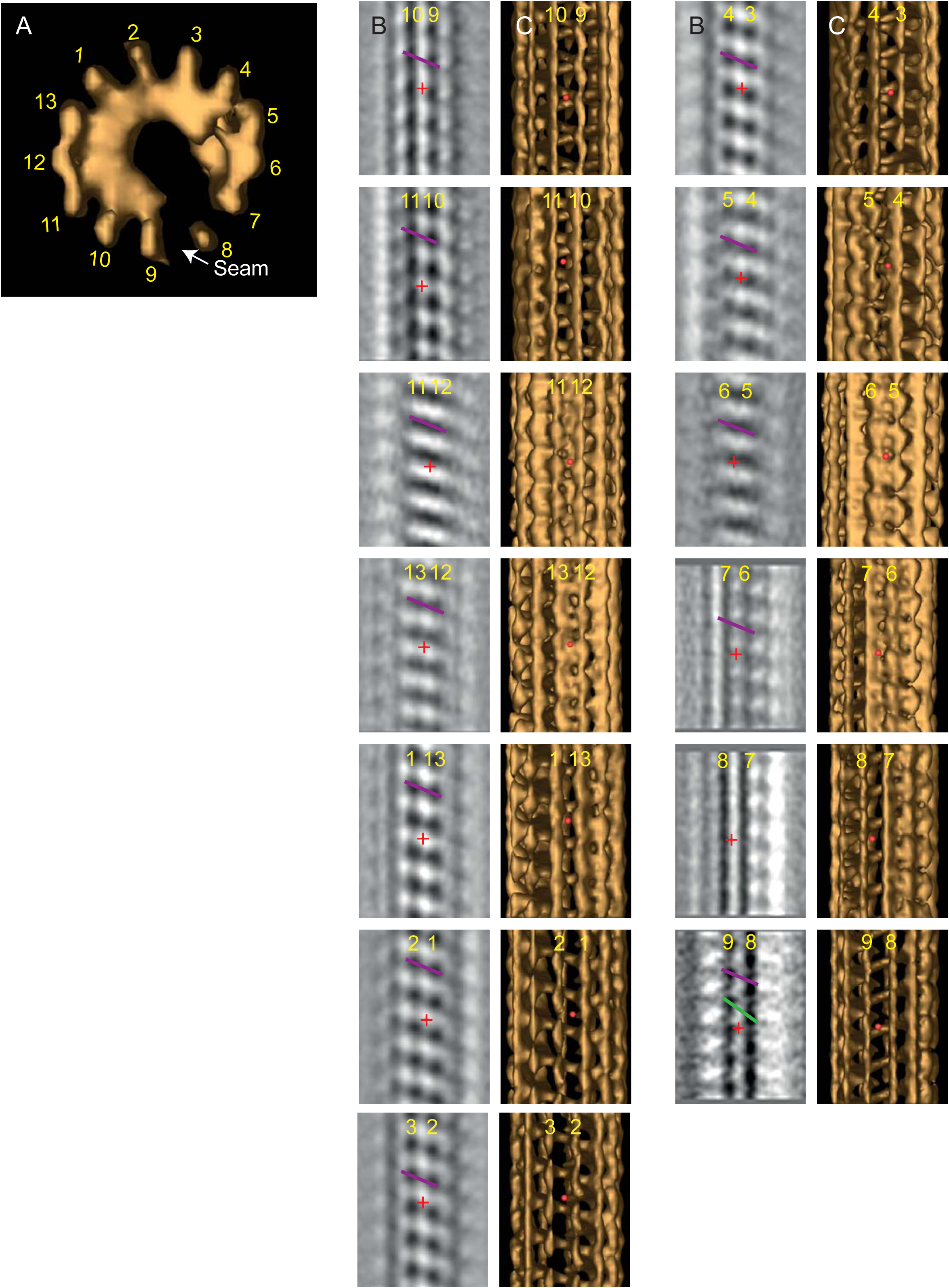
The location of the microtubule seam. A) A top-down view of the 3D model of the microtubule sub-tomogram average, including the TAILS helix. Protofilament numbers have been assigned based on the location of the seam in the dMT A-tubule as in Figure 4F and the numbers correlate to the images in B and C. B) Slices of the sub-tomogram average oriented such that two neighboring protofilaments are visualized. The red cross is the location of the red ball in C). The purple line shows the slope between neighboring tubulin subunits and fits all protofilament pairs except for pair 7-8 where no tubulin subunits could be identified, and pair 8-9 where the slope between protofilaments are different (green line) showing that the seam is here. C) A longitudinal view of the microtubule 3D model and the protofilaments.

## Supplementary movies

**Movie 1: The microtubule singlet region of the intact human spermatozoon.** All microtubules have a complete decoration of the TAILS complex. Each frame is a 0.7 nm thick slice from a cryo-electron tomogram. 7 frames/s. Scale bar 100 nm.

**Movie 2: Cross-sectional view of the microtubule sub-tomogram average and the electron densities that is the TAILS complex that rotates around the lumen.** 6 frames/s. Scale bar 10 nm.

**Movie 3: Longitudinal view of the microtubule sub-tomogram average containing the TAILS complex.** 6 frames/s. Scale bar 10 nm. 3D model of the sub-tomogram average showing the microtubule lattice in turquoise and the TAILS complex in green. 6 frames/s.

**Movie 4: The TAILS complex inside the doublet microtubule.** Scale bar 10 nm.

